# Disrupted neuropeptide-signaling drives enduring reward deficits after early-life stress

**DOI:** 10.64898/2026.08.26.747402

**Authors:** Matthew T. Birnie, Lara Taniguchi, Lia M. Harvey, Madison R. Tetzlaff, Lorenzo Mattioni, Amalia Floriou-Servou, Neeraj Thiagarajan, Graciella D. Angeles, Jennifer M. Daglian, Yuncai Chen, Tallie Z. Baram

**Affiliations:** Dept. of Pediatrics, University of California-Irvine, USA; Dept. of Anatomy & Neurobiology, University of California-Irvine, USA; Institute of Anatomy, University Medical Center of the Johannes Gutenberg-University Mainz, Germany; Dept. of Neurology, University of California-Irvine, USA

## Abstract

While brain systems mediating acute stress are essential for survival, chronic early-life stress (ELA) may lead to poor ability to experience pleasure (anhedonia), a core feature of depression. For decades, the stress neuropeptide corticotropin-releasing hormone (CRH) has been a target for treating depression, however the failure of several clinical trials blocking CRH receptor 1 (CRHR1) has left the therapeutic role of CRH signaling a major unresolved mystery. Here, we uncover the signaling plasticity behind this enigma with the use of in vivo G protein-coupled activation-based (GRAB) imaging and viral-genetic and pharmacological mechanistic manipulations. We find CRH signaling via CRHR1 indeed disrupts reward behaviors in naïve mice, but is dysregulated in ‘anhedonic’ mice with a history of ELA. Instead, activation of CRH receptor 2 (CRHR2) reverses anhedonia-like behaviors in adult ELA mice. These findings redefine our understanding of stress-mediated anhedonia and provide a precise, novel therapeutic target for stress-related mental illness.

## Introduction

Links between early-life stress (ELA) and adult affective disorders are well established ^1,2^, yet the underlying mechanisms remain unclear ^3–7^. ELA related problems such as depression and its core feature, anhedonia, are characterized by attenuated reward behaviors and augmented sensitivity to stress stimuli ^8–15^, suggesting that brain networks and molecules subserving these functions are enduringly impacted by ELA ^16–19^.

The evolutionarily conserved peptide, corticotropin releasing hormone (CRH), is canonically released from the hypothalamus to initiate the body’s response to stress ^20–22^, and elevated CRH levels in the cerebrospinal fluid and augmented CRH ‘drive’ have been hallmark correlates of depression ^23–25^. In addition to its role in the hypothalamic-pituitary-adrenal stress-response axis ^26^, CRH acts as a local neuromodulator in several brain circuits, including those involved in stress, threat and reward behaviors ^21,27–32^. The peptide and its receptors CRHR1 and CRHR2 are expressed in the amygdala, a brain region contributing to the processing of stressful stimuli, and in the nucleus accumbens (NAc), a hub of reward signaling ^33–38^. We recently identified a pathway between the basolateral amygdala (BLA) and the NAc expressing both CRH and GABA, and its stimulation provoked anhedonia-like behavior in naïve mice ^31,32,39^. Together, these observations in humans and experimental animals implicate CRH signaling in stress-related anhedonia and depression. Yet, despite this clear link, several well-designed clinical trials targeting the most abundant CRH receptor, CRHR1, have failed ^40–42^, leaving the potential therapeutic role of CRH signaling in depression a major unresolved mystery.

## Results

### CRH:CRHR1 signaling in the NAc regulates reward consumption and stress responses

To assess CRH:CRHR1 signaling in the naïve and ‘anhedonic’ NAc, we first confirmed that CRHR1 was robustly expressed on medium spiny neurons (MSNs) and interneuronal populations in the NAc (Figure 1A). Next, we determined the functional role of NAc CRHR1 signaling in reward behaviors germane to anhedonia using two distinct pharmacological receptor antagonists (Figure 1A - E). The home-cage reward consumption task utilizes the feeding experimental device (FED3 ^43^) on a free-feed, low motivation schedule that measures the number of sucrose pellets collected by a mouse. Blocking CRHR1 locally in the NAc with α-helicalCRH_9-41_, (a truncated form of CRH that binds to the receptor’s N-terminus but lacks the C-terminus domain to prevent activation and internalization ^44^), increased sucrose pellet consumption (Figure 1C, D). Similarly, a potent and highly selective small molecule allosteric inhibitor of CRHR1, NBI 30775 ^42^, augmented the consumption of sucrose pellets (Figure 1E, F).

**Figure 1:**
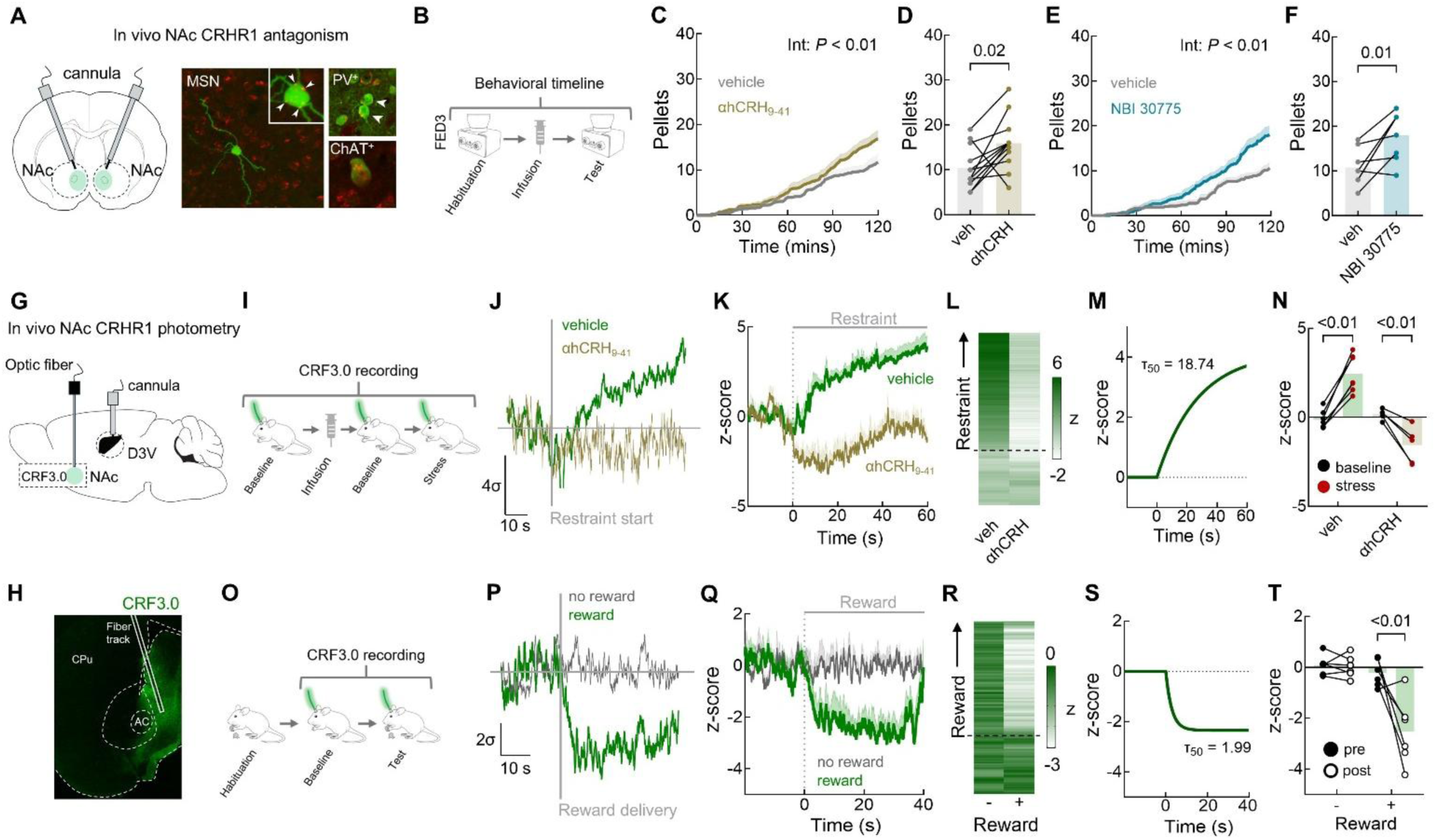
Opposing CRH release dynamics in the nucleus accumbens regulate reward consumption and stress responsiveness. **A**, Strategy for in vivo pharmacology targeting CRH receptor 1. **B**, Timeline of behavioral test and receptor manipulation. **C**, Consumption of sucrose pellets across a two-hour session with vehicle or α-helicalCRH_9-41_ infusions in the nucleus accumbens (two-way repeated measures analysis of variance (ANOVA) comparing drug x time, \**P* = 0.0043). *n* = 13 mice. **D**, Summary data of the total pellets consumed across the two-hour session (two-tailed paired *t*-test, \**P* = 0.02). *n* = 13 mice. **E**, Consumption of sucrose pellets across a two-hour session with vehicle or NBI 30775 infusions in the nucleus accumbens (two-way repeated measures analysis of variance (ANOVA) comparing drug x time, \**P* < 0.0001). *n* = 8 mice. **F**, Summary data of the total pellets consumed across the two-hour session (two-tailed paired *t*-test, \**P* = 0.01). *n* = 8 mice. **G**, Schematic diagram depicting the strategy for virus injection, fiber and cannula implantation, and measurement of CRF3.0 in the nucleus accumbens. **H**, Example CRF3.0 expression and fiber placement. **I**, **J**, Behavior stimulus and example signal in response to acute stress. **K**, **L**, Average traces of signal before and during a restraint stress. **M**, fitted one-phase association curve in response to restraint and on τ_50_. **N**, summary data of the change in signal measured before and during the restraint stress and following infusion of α-helicalCRH_9-41_ in the dorsal lateral ventricle (two-way repeated measures analysis of variance (ANOVA) comparing stress x drug, \**P* < 0.01, with Šídák’s post hoc test, veh: \**P* < 0.01; α-helicalCRH_9-41_: \**P* < 0.01,). *n* = 12 mice. **O**, **P**, Behavior stimulus and example signal in response to reward. **Q, R**, Average traces in signal before and during a reward presentation. **S**, fitted one-phase decay curve in response to reward and off τ_50_. **T**, summary data of the change in signal before and during a reward presentation (two-way repeated measures analysis of variance (ANOVA) comparing cue x reward, \**P* = 0.01, with Šídák’s post hoc test, \**P* < 0.01). *n* = 12 mice. Data represent mean ± SEM, and overlaid with individual data points where appropriate.

Next, to monitor *endogenous* CRH dynamics in response to reward and stress, we performed fiber photometry using the G-protein coupled receptor activation-based (GRAB) sensor CRF3.0 ^45^ (Figure 1G, H). We expressed the sensor in the medial NAc, a known hedonic hotspot and the termination site of CRH-expressing BLA-origin projection neurons ^31,32,39,46^. Because CRH is released by stress in several brain regions ^47^, we first used immobilization stress to validate the sensor’s sensitivity and specificity, by measuring the signal in response to an acute immobilization stressor (60 s; Figure 1I). As expected, this aversive stimulus elicited a sharp increase in signal that persisted throughout the stress period and was abolished by intracerebroventricular (i.c.v) infusion of α-helicalCRH_9-41_, confirming the sensor’s fidelity for detecting peptide release and signaling via CRHR1 (Figure 1J - N). We next tested CRH release in response to receipt of sucrose pellet rewards. As opposed to the effects of stress, the delivery of these pellets induced a robust and sustained *reduction* of CRH fluorescence (Figure 1P - T). Overall, these combined photometry and pharmacological data show that NAc CRHR1 signaling gates stress and reward behaviors.

### NAc CRH:CRHR1 signaling promotes anhedonia-like behavior

Our data suggested that NAc CRH:CRHR1 signaling mediates physiological responses to acute stress and gates the response to rewarding stimuli. To test this supposition, we augmented endogenous NAc CRH levels using both pharmacological and viral-genetic approaches. We first leveraged the fact that the CRH binding protein (CRH-BP) acts to sequester CRH away from its receptors ^48,49^. Indeed, in humans, ∼40% of cortical CRH is bound to the CRH-BP, and this protein is expressed in mouse NAc (Figure 2A). Therefore, we infused the CRH-BP blocker CRH_6-33_ that displaces endogenous CRH from CRH-BP and that does not act on the CRHR1 receptor, ^50^ directly into the medial NAc (Figure 2B). Displacing endogenous CRH from the binding protein significantly reduced pellet consumption in the FED3 sucrose pellet reward task described above (Figure 2C, D). The efficacy of this manipulation in increasing ambient CRH levels in the NAc was validated by recording CRF3.0 fluorescence during CRH_6-33_ infusion (Figure 2E). Consistent with its expected function, CRH_6-33_ elicited a robust increase in sensor fluorescence (Figure 2F - I), confirming that elevated NAc CRH levels directly correlated with impaired reward behaviors.

**Figure 2:**
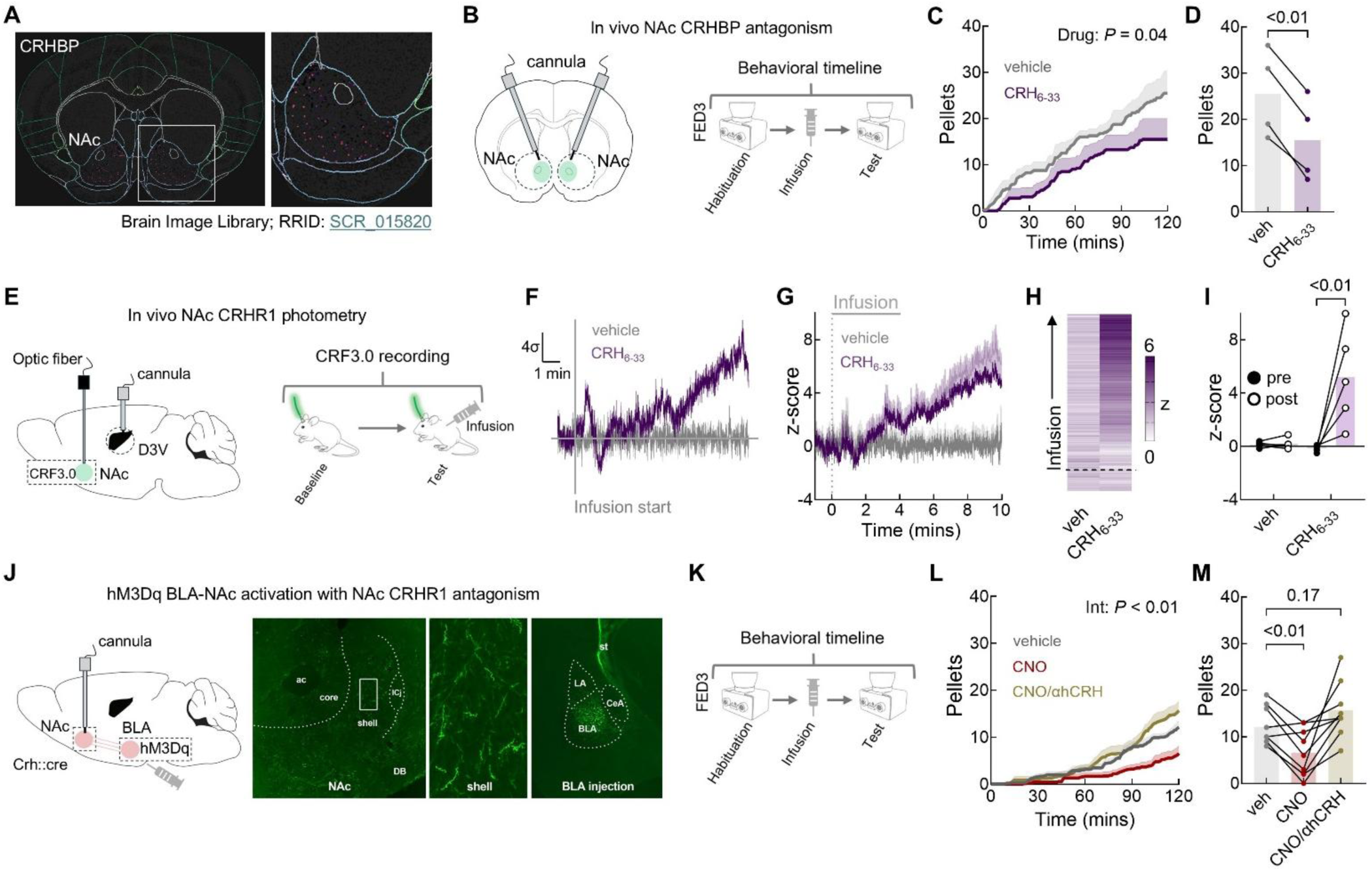
CRHR1 signaling in the nucleus accumbens negatively regulates reward behaviors. **A,** Expression of CRH-binding protein in the nucleus accumbens**. B**, Strategy for in vivo pharmacology and behavioral timeline. **C**, Consumption of sucrose pellets across a two-hour session with vehicle or CRH_6-33_ infusions, (two-way repeated measures analysis of variance (ANOVA) comparing drug, \**P* = 0.0421). *n* = 4 mice. **D**, Summary data of the total pellets consumed across the two-hour session (two-tailed paired *t*-test, \**P* < 0.01). *n* = 4 mice. **E**, Combined in vivo CRF3.0 fiber photometry and pharmacology. **F**, Example signal in response to CRH_6-33_ infusion. **G**, **H**, Average traces in CRF3.0 signal before, during and after CRH_6-33_ infusion. **I**, Summary data of the change in signal before and after infusion (two-way repeated measures analysis of variance (ANOVA) comparing drug x time, \**P* = 0.01, with Šídák’s post hoc test, \**P* < 0.01). *n* = 10 mice. **J**, Strategy for combining hM3Dq activation of a basolateral amygdala-nucleus accumbens CRH+ projection and CRHR1 antagonism. **K**, Behavior stimulus. **L**, Consumption of sucrose pellets across a two-hour session with vehicle, CNO, or CNO and α-helicalCRH_9-41_ infusions (two-way repeated measures analysis of variance (ANOVA) comparing drug x time, \**P* = 0.0052). *n* = 9 mice. **M**, Summary data of the total pellets consumed across the two-hour session (one-way repeated measures analysis of variance (ANOVA) with Šídák’s post hoc test, \**P* < 0.01, comparing vehicle versus CNO, and vehicle versus α-helicalCRH_9-41_). *n* = 9 mice. Data represent mean ± SEM, and overlaid with individual data points where appropriate.

In addition to increasing NAc CRH expression by locally displacing the peptide from its binding protein, we stimulated CRH release from a long-range input using a chemogenetic approach. Building on our prior discovery of a CRH / GABA-expressing projection from BLA to NAc ^31,32,39^, we virally expressed Cre-dependent hM3Dq DREADDs in the BLA of CRH-Cre mice and delivered clozapine-n-oxide (CNO) locally to axon terminals in the NAc (Figure 2J, K). Stimulating this CRH^+^ BLA-NAc projection significantly decreased sucrose consumption (Figure 2L, M). However, because these long-range neurons can release both CRH and GABA, we sought to isolate the role of CRH in this reward task. To this end, we infused into the NAc the CRHR1 antagonist α-helicalCRH_9-41_ in combination with CNO. Strikingly, α-helicalCRH_9-41_ successfully blocked the CNO-induced suppression of reward behavior (Figure 2L, M), establishing that the observed anhedonia-like behavior is driven by CRH:CRHR1 signaling rather than GABAergic transmission in the NAc.

Together, these data demonstrate that elevating NAc CRH levels, either through local displacement from the CRH-BP, or by stimulating a long-range CRH^+^ BLA-NAc projection is sufficient to suppress reward consumption, buttressing the role of CRH and CRH:CRHR1 signaling in gating reward behaviors.

### Early-life stress promotes maladaptive CRH:CRHR1 signaling to drive reward deficits

Having established that the NAc CRH:CRHR1 signaling modulates reward behavior in naive mice (Figures 1, 2), we next investigated whether early-life stress (ELA) dysregulates this signaling. To recapitulate the human link between ELA and anhedonia and depression, we employed the translationally relevant limited bedding and nesting (LBN) paradigm of ELA ^51–53^ (Figure 3A), which reliably generates dysregulated adult reward behaviors ^30–32^. Using this model, we found that ELA significantly decreased sucrose consumption in males (Figure 3B, C) and increased motivation for palatable food in females (Figure S1). Notably, these reward-related deficits were independent of any potential changes in anxiety-like behavior or sensorimotor function (Figure S2).

**Figure 3:**
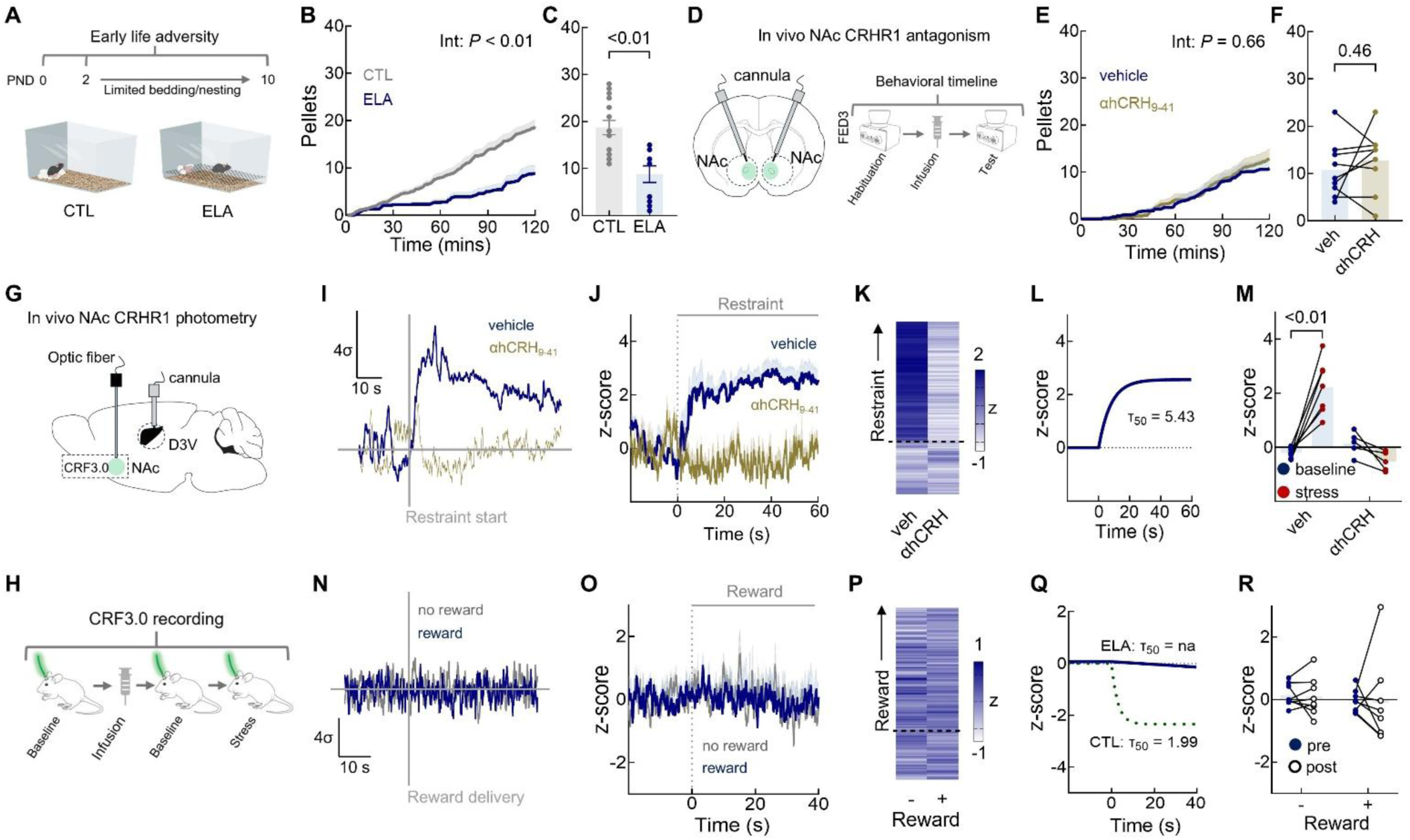
Early-life stress dysregulates NAc CRHR1 signaling. **A**, Timing of limited bedding and nesting model of early-life stress (ELA). **B**, Consumption of sucrose pellets in adult control and ELA mice across a two-hour session (two-way repeated measures analysis of variance (ANOVA) comparing time x experience, \**P* = 0.0001). *n* = 25 mice. **C**, Summary data of the total pellets consumed during a two-hour session (two-tailed paired *t*-test, \**P* < 0.01). *n* = 25 mice. **D**, Strategy for in vivo pharmacology. **E**, Consumption of sucrose pellets over a two-hour session following vehicle or α-helicalCRH_9-41_ infusions into the nucleus accumbens (two-way repeated measures analysis of variance (ANOVA) comparing drug x time, *P* = 0.6598). *n* = 9 mice. **F**, Summary data of the total pellets consumed across the two-hour session (two-tailed paired *t*-test, *P* = 0.42). *n* = 9 mice. **G**, **H**, Combined in vivo CRF3.0 fiber photometry and pharmacology and behavioral procedure. **I**, Example signal in response to acute stress. **J**, **K**, Average traces of signal before and during a restraint stress. **L**, fitted one-phase association curve in response to restraint and on τ_50_. **M**, Summary data of the change in signal measured before and during the restraint stress and following infusion of α-helicalCRH_9-41_ in the dorsal lateral ventricle (two-way repeated measures analysis of variance (ANOVA) comparing stress x drug, \**P* < 0.01, with Šídák’s post hoc test, \**P* < 0.01). *n* = 12 mice. **N**, Example signal in response to a reward cue. **O**, **P**, Average traces in signal. **Q**, fitted one-phase decay curve in response to reward and off τ_50_. **R**, Summary data of the change in signal before and during reward pellet presentation (two-way repeated measures analysis of variance (ANOVA) comparing cue x reward, *P* = 0.5541). *n* = 16 mice. Data represent mean ± SEM, and overlaid with individual data points where appropriate.

To determine if NAc CRH:CRHR1 signaling is responsible for reward-related behavioral deficits after ELA, we combined pharmacological approaches with behavioral measures. Because CRHR1 blockers have been used as therapeutic agents for depression ^40–42^, we first assessed whether intra-NAc infusion of α-helicalCRH_9-41_ could rescue reward behavior in ‘anhedonic-like’ ELA mice (Figure 3D - F). Strikingly, the antagonist failed to increase sucrose consumption in ELA mice (Figure 3E, F), a result that stands in stark contrast to its efficacy in control animals (Figure 1). This divergence suggests that ELA induces an enduring shift in CRHR1 function or availability.

We first excluded the possibility that the lack of efficacy of the CRHR1 blocker is a result of major reduction of CRHR1 expression in adult ‘anhedonic’ ELA mice (Figure S3). Next, to directly examine whether ELA-induced adult anhedonia involves altered CRH:CRHR1 dynamics, we combined fiber photometry with pharmacological blockade (Figure 3G - R). Similar to naïve control mice, acute stress generated a robust fluorescence signal increase in ELA mice (Figure 3I - M; Figure S4). However, in contrast to control mice (Figure 1P - T), reward cues failed to inhibit NAc CRH signaling in anhedonic-like ELA mice (Figure 3N - R). These results indicate that ELA selectively disrupts the reward-related inhibitory action of the CRH:CRHR1 signaling while leaving acute stress-evoked activation intact.

We reasoned that ELA may dysregulate reward behaviors by generating enduringly receptor-saturating levels of CRH within the NAc. To test this possibility, we determined whether artificially increasing endogenous CRH in the NAc influences physiological and behavioral responses in a manner observed in controls (Figure 2). Using the same methodological approaches described for naïve mice, we found that local displacement of CRH from its binding protein failed to reduce sucrose consumption further in ‘anhedonic’ ELA mice (Figure S5A - C). Similarly, chemogenetic stimulation of the CRH^+^ BLA-NAc projection did not further reduce palatable food consumption (Figure S5D - F). The inability to further suppress reward behaviors is consistent with high, receptor-saturating NAc CRH levels in ELA mice. To test this possibility directly, we combined fiber photometry and pharmacology to measure CRH:CRHR1 binding. Indeed, CRH displacement via CRH_6-33_ infusion failed to elicit an increase in sensor fluorescence (Figure S5G - K), in contrast with the robust signal seen in naïve, control mice (Figure 2).

Together, these data demonstrate that ELA leads to chronic CRH:CRHR1 signaling in the adult NAc, which drives long-term suppression of reward behaviors.

### CRHR2 activation selectively rescues reward deficits induced by ELA

CRH modulates signaling in the brain primarily via its high-affinity receptor, CRHR1^33^. Accordingly, the above experiments demonstrate that CRH release in the NAc, originating from both local and long-range sources, suppresses reward consumption in control mice by acting on CRHR1 (Figures 1, 2). However, in the context of anhedonia, augmenting endogenous CRH in the NAc with DREADDs or binding-protein displacement did not further suppress reward behaviors (Figure S5). Importantly, blocking CRHR1 failed to rescue these behaviors (Figure 3G, H; Figure S1), recapitulating the failure of CRHR1 blockers in clinical trials for depression ^40,41^.

Notably, in addition to the high-affinity CRHR1, the CRH system includes the lower-affinity CRHR2. While CRHR2 has primarily been studied for its peripheral role in homeostatically regulating cardiovascular, inflammatory, and metabolic processes ^54–56^, it is also expressed in the brain, where it contributes to stress response regulation ^57,58^, often via urocortin 2 and 3 ^59^ but also via CRH ^60,61^.Therefore, we tested whether NAc CRHR2 could regulate reward behavior in ‘anhedonic’ ELA mice.

First, we determined the spatial and cellular distribution of the CRHR2 in the NAc, together with that of CRH, CRHR1 and CRH-BP using Multiplexed Error-Robust Fluorescence in situ Hybridization (MERFISH) ^62^. We clustered cell populations using established markers for medium spiny neurons (*Drd1*, *Drd2*), interneurons (*Sst*, *Chat*), and glia (Figure 4A, B) and identified robust expression of *Crh*, *Crhr1*, *Crhr2*, and *Crhbp*. Specifically, *Crh* and *Crhr2* were localized to *Drd1^+^* and *Drd2^+^* neurons, while *Crhr1* was expressed in in *Drd1^+^*, *Drd2^+^* as well as in interneuronal *Chat^+^* populations. *Crhbp* expression was restricted to *Sst^+^* interneurons (Figure 4A - D; Figure S6), as described for the human NAc ^63^.

**Figure 4:**
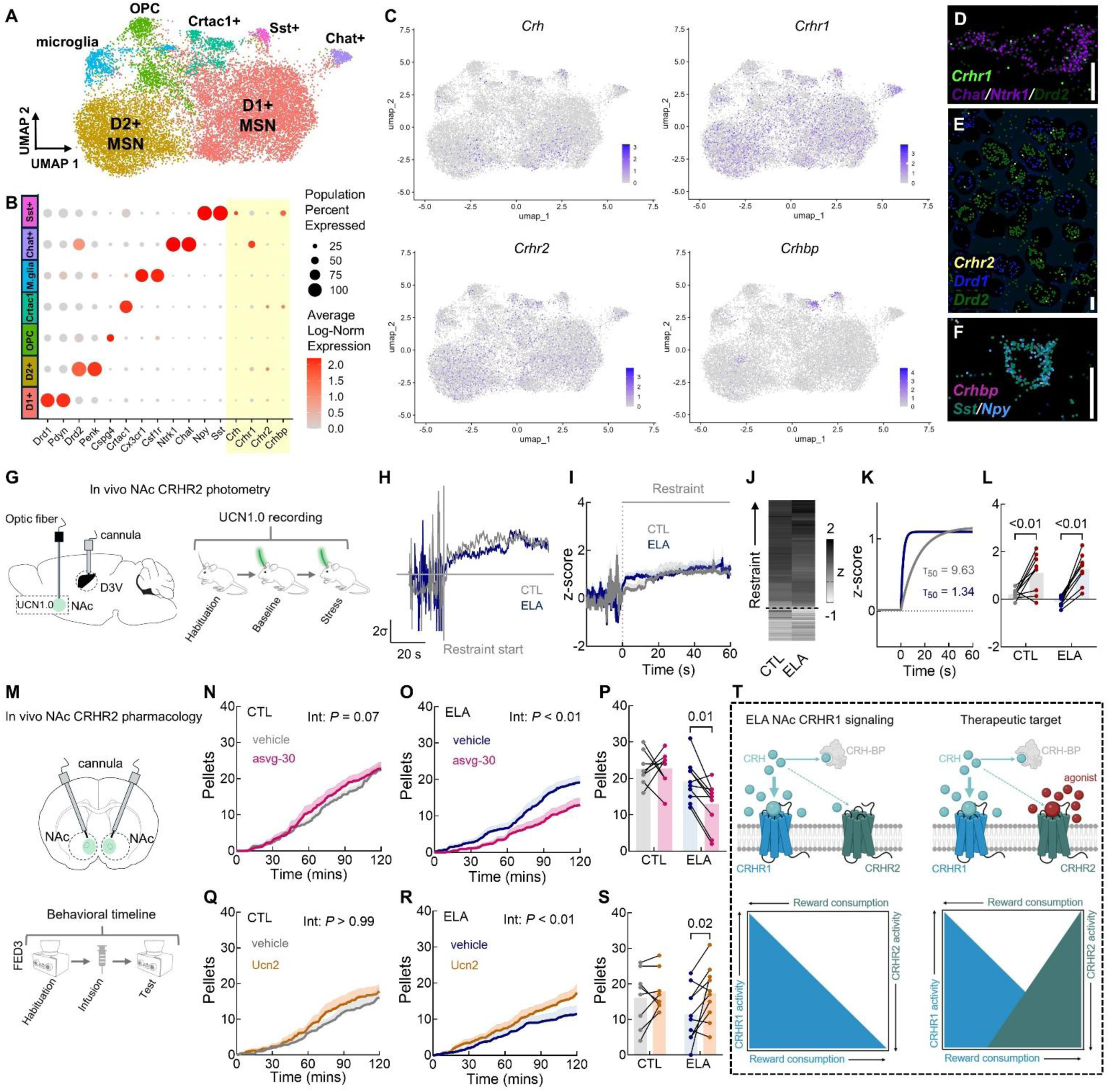
NAc CRHR2 signaling mediates restored reward behaviors in ‘anhedonic’ ELA mice. **A**, Uniform manifold approximation and projection (UMAP) of the MERFISH data with cell clusters colored by cell type. **B**, Gene expression data demonstrates top cell cluster markers. **C**, UMAPs of *Crh*, *Crhr1*, *Crhr2* and *Crhbp*. **D** - **F**, Representative MERscope images of *Crhr1*, *Crhr2* and *Crhbp* expression in dopamine receptor D2-positive (*Drd2*) medium spiny neurons, and choline acetyltransferase-positive (*Chat*) and somatostatin-positive (*Sst*) interneurons. **G**, Combined in vivo UCN1.0 fiber photometry and pharmacology. **H**, Example signal in response to acute stress. **I**, **J**, Average traces of signal before and during a restraint stress. **K**, fitted one-phase association curve in response to restraint and on τ_50_. **L**, summary data of the change in signal measured before and during the restraint stress in control and ELA mice (two-way repeated measures analysis of variance (ANOVA) comparing stress, \**P* < 0.01, with Šídák’s post hoc test, \**P* < 0.01). *n* = 18 mice. **M**, Strategy for in vivo pharmacology and behavioral procedure. **N**, Consumption of sucrose pellets in control mice across a two-hour session following vehicle or antisauvagine-30 infusions in the nucleus accumbens (two-way repeated measures analysis of variance (ANOVA) comparing time x drug, *P* = 0.07). *n* = 8 mice. **O**, In ELA mice (two-way repeated measures analysis of variance (ANOVA) comparing time x drug, \**P* < 0.0001). *n* = 9 mice. **P**, Summary data of the total pellets consumed across a two-hour session (two-way repeated measures analysis of variance (ANOVA) comparing drug x experience, \**P* = 0.045, with Šídák’s post hoc test, \**P* = 0.011). *n* = 18 mice. **Q**, Consumption of sucrose pellets in control mice across a two-hour session following vehicle or urocortin 2 infusions in the nucleus accumbens (two-way repeated measures analysis of variance (ANOVA) comparing time x drug, *P* = 0.9986). *n* = 8 mice. **R**, In ELA mice (two-way repeated measures analysis of variance (ANOVA) comparing time x drug, *P* < 0.0001). *n* = 10 mice. **S**, Summary data of the total pellets consumed across the two-hour session (two-way repeated measures analysis of variance (ANOVA) comparing drug, \**P* = 0.02, with Šídák’s post hoc test, \**P* = 0.02). *n* = 18 mice. **T**, Model proposing the therapeutic mechanism for activating CRHR2. Following ELA, increasing CRH:CRHR1 activity decreases reward consumption. Thus, to equilibrate CRH:CRHR1 activity, stimulating CRHR2 with selective modulators can increase reward consumption to normalize behavior. Data represent mean ± SEM, and overlaid with individual data points where appropriate.

Leveraging these findings, we next investigated the ability of CRHR2 activation to modulate reward behaviors. We first employed fiber photometry to monitor endogenous CRHR2 activity dynamics using the GRAB sensor UCN1.0, which is based on a modified CRHR2 (Figure 4G) ^45^. In both control and anhedonic ELA mice, acute immobilization stress elicited a significant increase in fluorescence above baseline (Figure 4H - L). This robust signaling indicates that, unlike the reward-related CRH signaling derangements, the fundamental functions of CRHR1 and CRHR2 in modulating response to stress remain intact following ELA (Figure S4).

Next, we used pharmacological manipulations to determine if CRHR2 signaling influenced reward-related behavior. Blocked CRHR2 activity via direct infusion of antisauvagine-30, a selective competitive antagonist ^64^ (Figure 4M - P) had no effect in control mice (Figure 4N, P). By contrast, in adult ELA mice, antisauvagine-30 further exacerbated the deficit in sucrose consumption (Figure 4O, P). These data support the scenario that the lower-affinity CRHR2 exerts little function in control mice, in which NAc CRH levels are relatively low. In contrast, the receptor serves as a homeostatic brake during high CRH levels, such as in anhedonic adult ELA mice. To empirically test this possibility, we stimulated CRHR2 using the selective agonist Urocortin 2 in both control and ‘anhedonic’ ELA mice. While Urocortin 2 had no effect in controls (Figure 4Q, S), it strikingly rescued reward deficits in ELA mice (Figure 4R, S), restoring their palatable food consumption to control levels.

Collectively, these data indicate that selective activation of CRHR2 in anhedonic mice potently counteracts the enduring, maladaptive CRHR1 signaling induced by early-life stress, effectively restoring reward behaviors (Figure 4T).

While the current studies establish a direct role for CRH signaling in reward behavior, the peptides function in a complex in vivo milieu includes additional neurotransmitters (glutamate and GABA), and neuromodulators (e.g., dopamine, oxytocin, PACAP). CRH:CRHR1 signaling regulates the release of dopamine and oxytocin (and potentially others) in the NAc ^34,35,65^, such that aberrant CRH:CRHR1 signaling may have far-ranging implication for a variety of motivated behaviors ^66–68^. Future studies combining multi-neurotransmitter and -neuropeptide-specific sensors with in vivo CRISPR-Cas9 gene manipulations of the CRHRs would shed further light on the direct and indirect roles of CRH-related targets in reward-related behaviors in health and disease.

## Conclusions

In summary, we demonstrate that CRH signaling in the NAc is a critical modulator of reward behaviors and is permanently impacted by early-life stress, leading to reward deficits germane to human anhedonia and depression. We identify the underlying mechanism as disrupted signaling of CRH through its primary high affinity receptor, CRHR1, potentially as a result of chronic receptor saturation. Our discovery of ELA-induced dysregulation of CRH:CRHR1 signaling in anhedonic mice identifies a mechanistic cause for the lack of success of CRHR1 blockers in clinical trials for neuropsychiatric disorders ^40,41^. Importantly, we offer an alternative neurobiologically grounded target for intervention: by showing that enhancing the activity of the less well-studied and lower affinity CRH receptor, CRHR2, robustly rescues reward behaviors in anhedonic mice with a history of ELA.

## Acknowledgements

We thank the Yulong Li laboratory for sharing their expertise and information on the selectivity and specificity of their CRHR1 (AAV-hSyn-CRF3.0) and CRHR2 (AAV-hSyn-UCN1.0) GRAB sensors.

## Author contributions

Conceptualization: TZB, MTB

Methodology: MTB, MRT, AFS, YC, TZB

Investigation: MTB, LT, LH, MRT, LM, AFS, NT, GDA, JMD, YC

Visualization: MTB, MRT, YC

Funding acquisition: TZB, MTB

Project administration: TZB Supervision: TZB, MTB

Writing – original draft: TZB, MTB

Writing – review & editing: MTB, MRT, YC,TZB

## Data, code, and materials availability

Source data are provided with this paper. Additional data that supports the findings of this study are available from the corresponding author upon request.

## Supplementary Materials

Materials and Methods

Figures S1 – 6

Table S1

---

**Figure S1:**
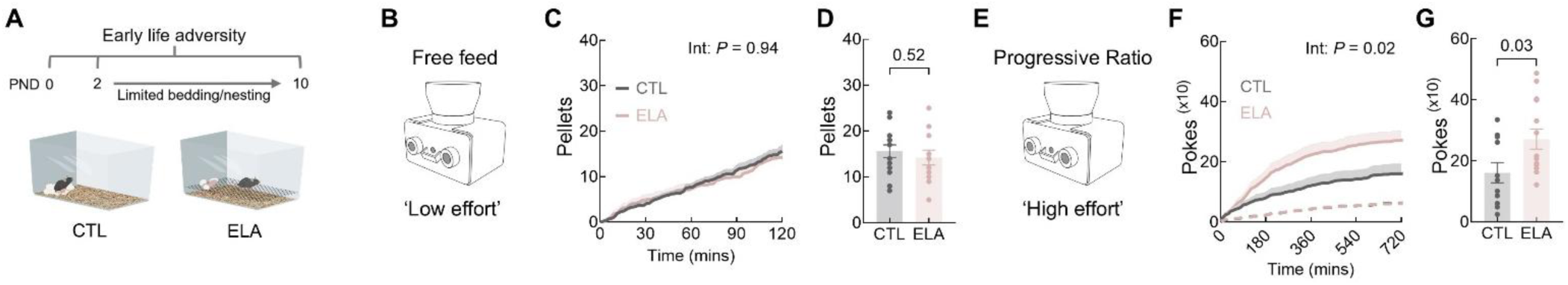
Low and high effort reward behaviors in adult control and ELA females. **A**, Timing of limited bedding and nesting model of early-life stress. **B**, Behavior paradigm – low effort. **C**, Consumption of sucrose pellets in control and early-life stress mice across a two-hour session under free conditions (two-way repeated measures analysis of variance (ANOVA) comparing time x experience, *P* = 0.9377). *n* = 28 mice. **D**, Summary data of the total pellets consumed a two-hour session (two-tailed unpaired *t*-test, *P* = 0.52). *n* = 25 mice. **E**, Behavior paradigm – high effort. **F**, Consumption of sucrose pellets in control and early-life stress mice across a two-hour session under high motivation – progressive ratio (two-way repeated measures analysis of variance (ANOVA) comparing time x experience, \**P* = 0.017). *n* = 25 mice. **G**, Summary data of the total pellets consumed a two-hour session (two-tailed paired *t*-test, \**P* = 0.03). *n* = 25 mice. Data represent mean ± SEM, and overlaid with individual data points where appropriate.

**Figure S2:**
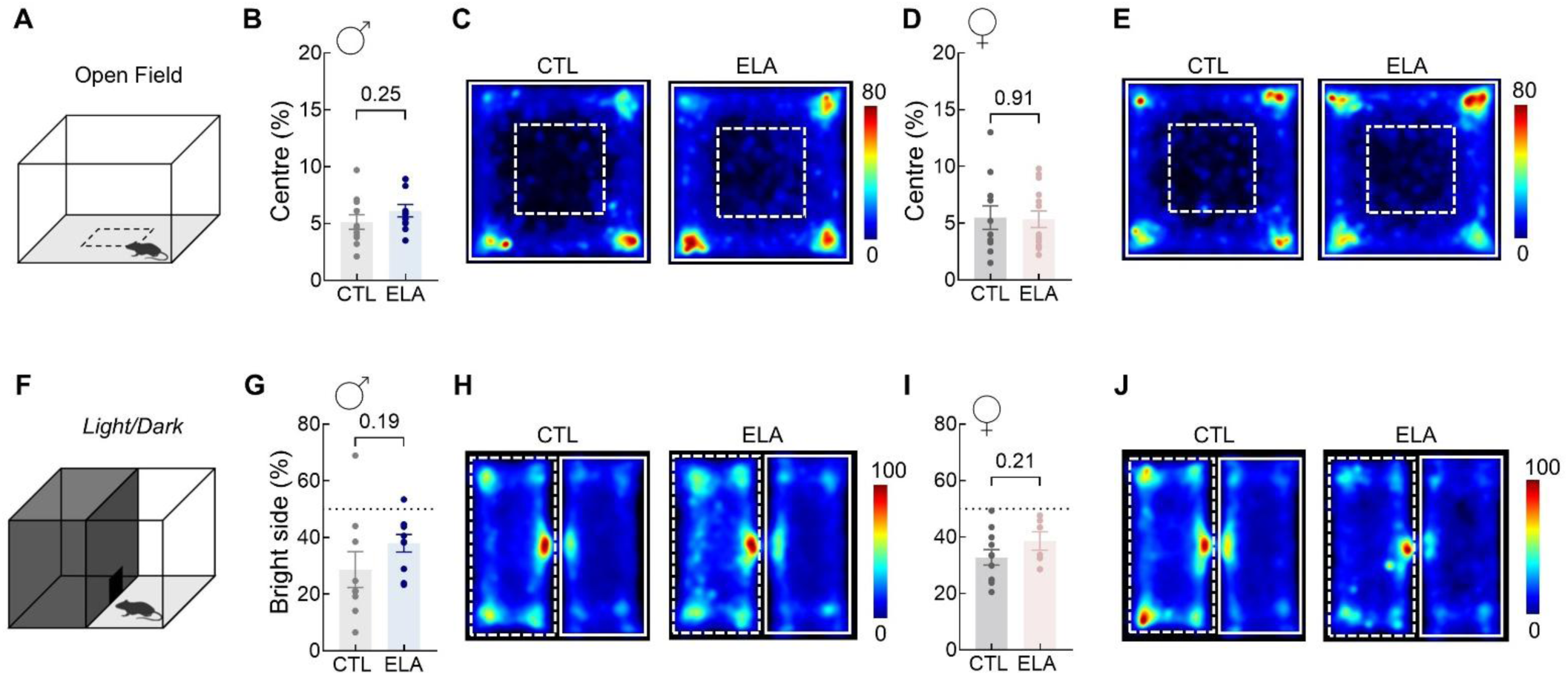
Assessment of anxiety-like and sensorimotor performance in control and ELA mice. **A**, cartoon depicting the setup of the open field behavior test. **B**, Percent of time control and ELA males spent in the center of an open field (two-tailed unpaired *t*-test, *P* = 0.25). *n* = 22 mice. **C**, Heatmap visualization of open field occupancy. **D**, Percent of time control and ELA females spent in the center of an open field (two-tailed unpaired *t*-test, *P* = 0.91). *n* = 25 mice. **E**, Heatmap visualization of open field occupancy. **F**, Behavior test. **G**, Percent of time control and ELA males spent in the brightly lit chamber (two-tailed unpaired *t*-test, *P* = 0.19). *n* = 19 mice. **H**, Heatmap visualization of bright chamber (solid white box) occupancy. **I**, Percent of time control and ELA females spent in the brightly lit chamber (two-tailed unpaired *t*-test, *P* = 0.21). *n* = 17 mice. **J**, Heatmap visualization of bright chamber (solid white box) occupancy.

**Figure S3:**
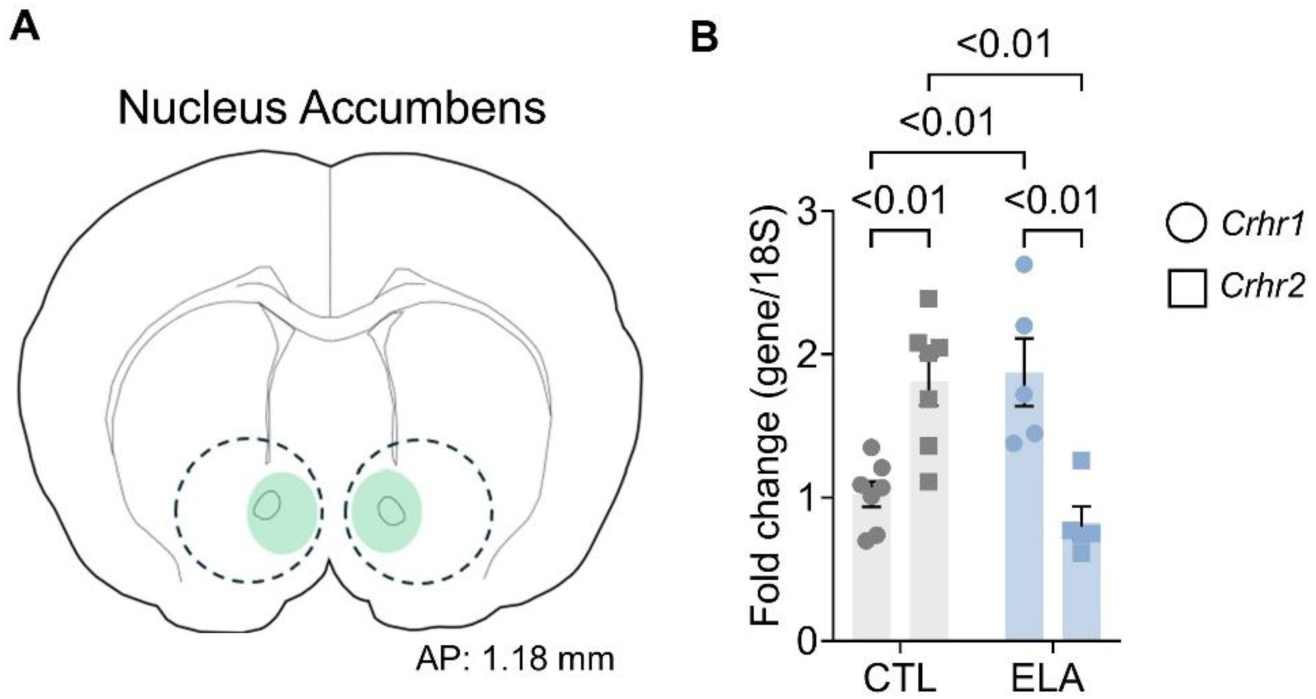
Expression levels of CRHR1 and CRHR2 in the nucleus accumbens of adult mice raised in control or ELA conditions. **A**, Location of nucleus accumbens micropunch tissue (green) from CTL and ELA mice. **B**, mRNA expression of CRHR1 and CRHR2 normalizing expression to control CRHR1 (two-way ANOVA comparing gene x experience, P < 0.01). *n* = 12 mice.

**Figure S4:**
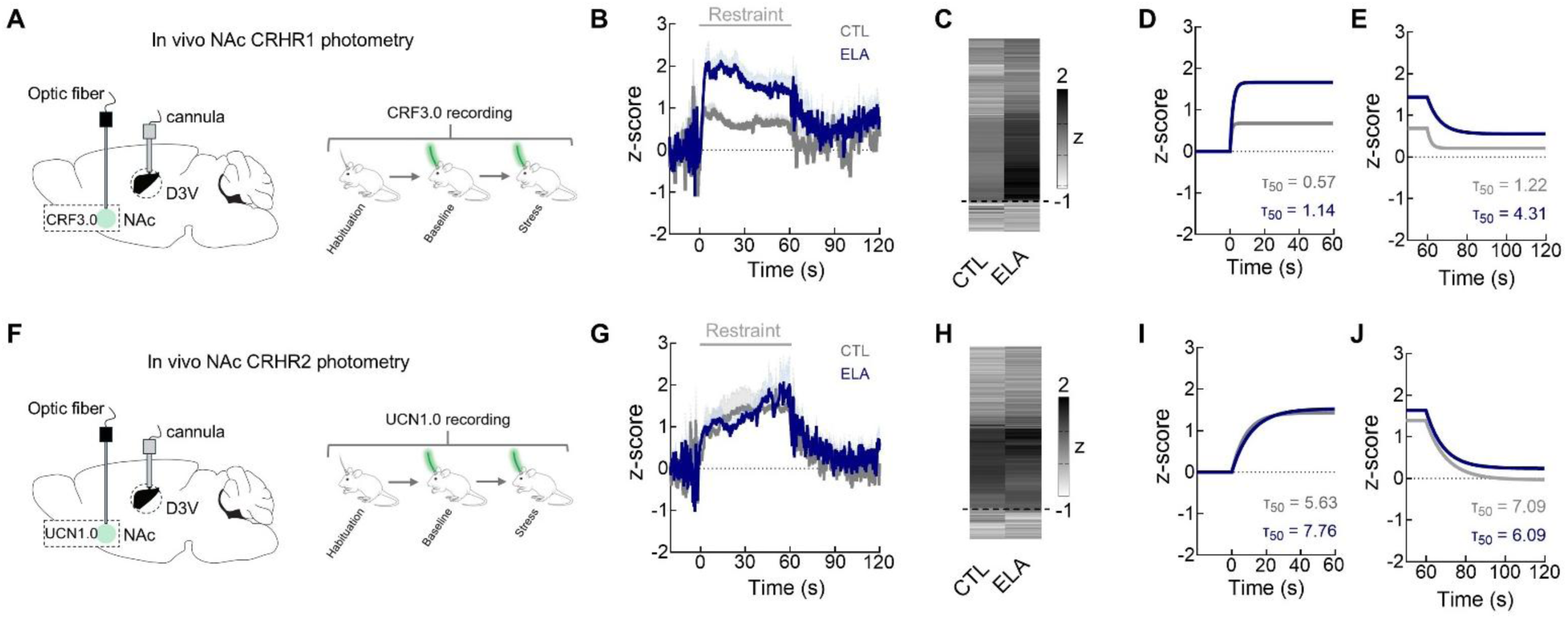
Comparative CRF3.0 and UCN1.0 signaling dynamics in control and ELA mice in response to acute stress. **A**, Combined in vivo CRF3.0 fiber photometry. **B**, **C**, Average traces in signal before, during and after acute stress. **D**, **E**, fitted curves of on (d) and off (e) τ_50_ of the change in fluorescence. *n* = 16 mice. **F**, Combined in vivo UCN1.0 fiber photometry. **G**, **H**, Average traces in signal before, during and after acute stress. **I**, **J**, fitted curves of on (d) and off (e) τ_50_ of the change in fluorescence. *n* = 20 mice.

**Figure S5:**
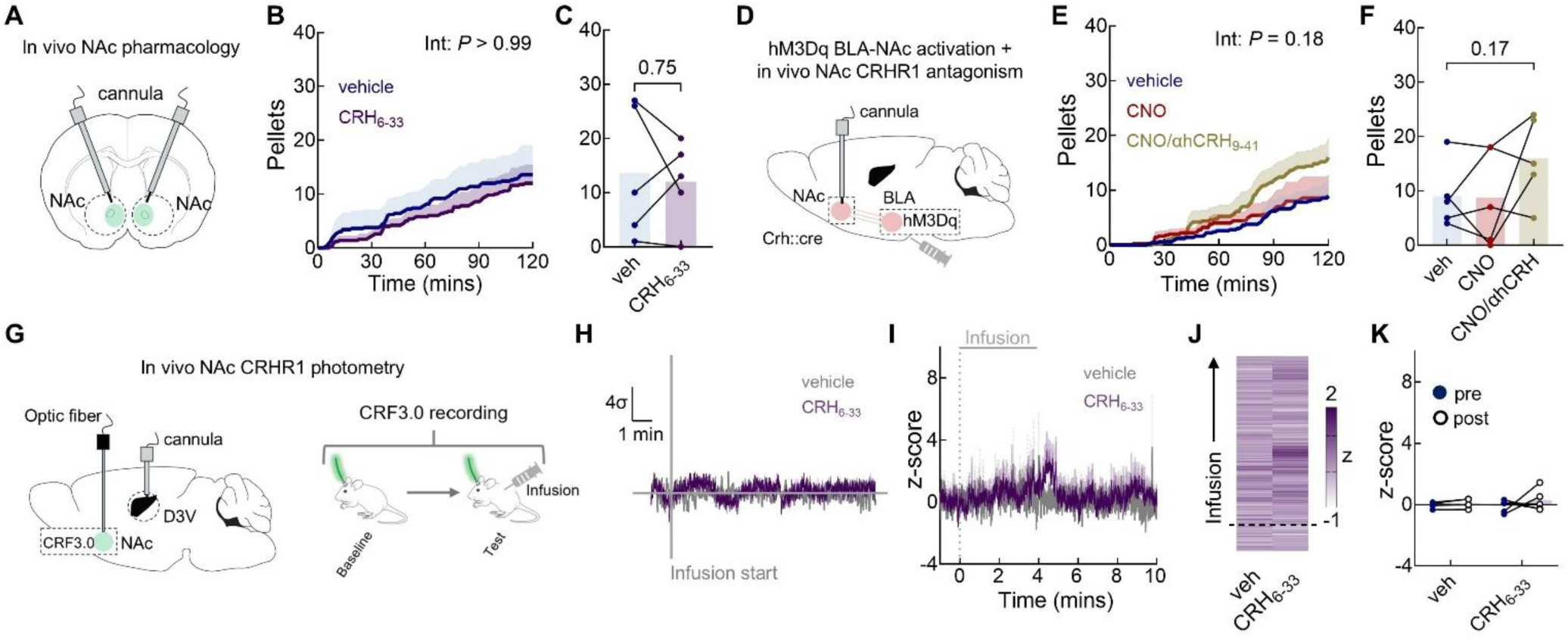
ELA-induced dysregulation of CRHR1 signaling renders the NAc unresponsive to further elevation of peptide levels. **A**, In vivo pharmacology. **B**, Consumption of sucrose pellets across a two-hour session with vehicle or CRH_6-33_ infusions, (two-way repeated measures analysis of variance (ANOVA) comparing time x drug, *P* < 0.9999). *n* = 5 mice. **C**, Summary data of the total pellets consumed across the two-hour session (two-tailed paired *t*-test, *P* = 0.75). *n* = 5 mice. **D**, Strategy for combining hM3Dq activation of a basolateral amygdala-nucleus accumbens CRH+ projection and CRHR1 antagonism. **E**, Consumption of sucrose pellets across a two-hour session with vehicle, CNO, or CNO and α-helicalCRH_9-41_ infusions (two-way repeated measures analysis of variance (ANOVA) comparing drug x time, *P* = 0.1758). *n* = 5 mice. **F**, Summary data of the total pellets consumed across the two-hour session (one-way repeated measures analysis of variance (ANOVA)), *P* = 0.1739. **G**, Combined in vivo CRF3.0 fiber photometry and pharmacology. **H**, Example signal in response to CRH_6-33_ infusion. **I**, **J**, Average traces in CRF3.0 signal before, during and after CRH_6-33_ infusion. **K**, Summary data of the change in signal before and after infusion (two-way repeated measures analysis of variance (ANOVA) comparing drug x time, *P* = 0.55). *n* = 10 mice. Data represent mean ± SEM, and overlaid with individual data points where appropriate.

**Figure S6:**
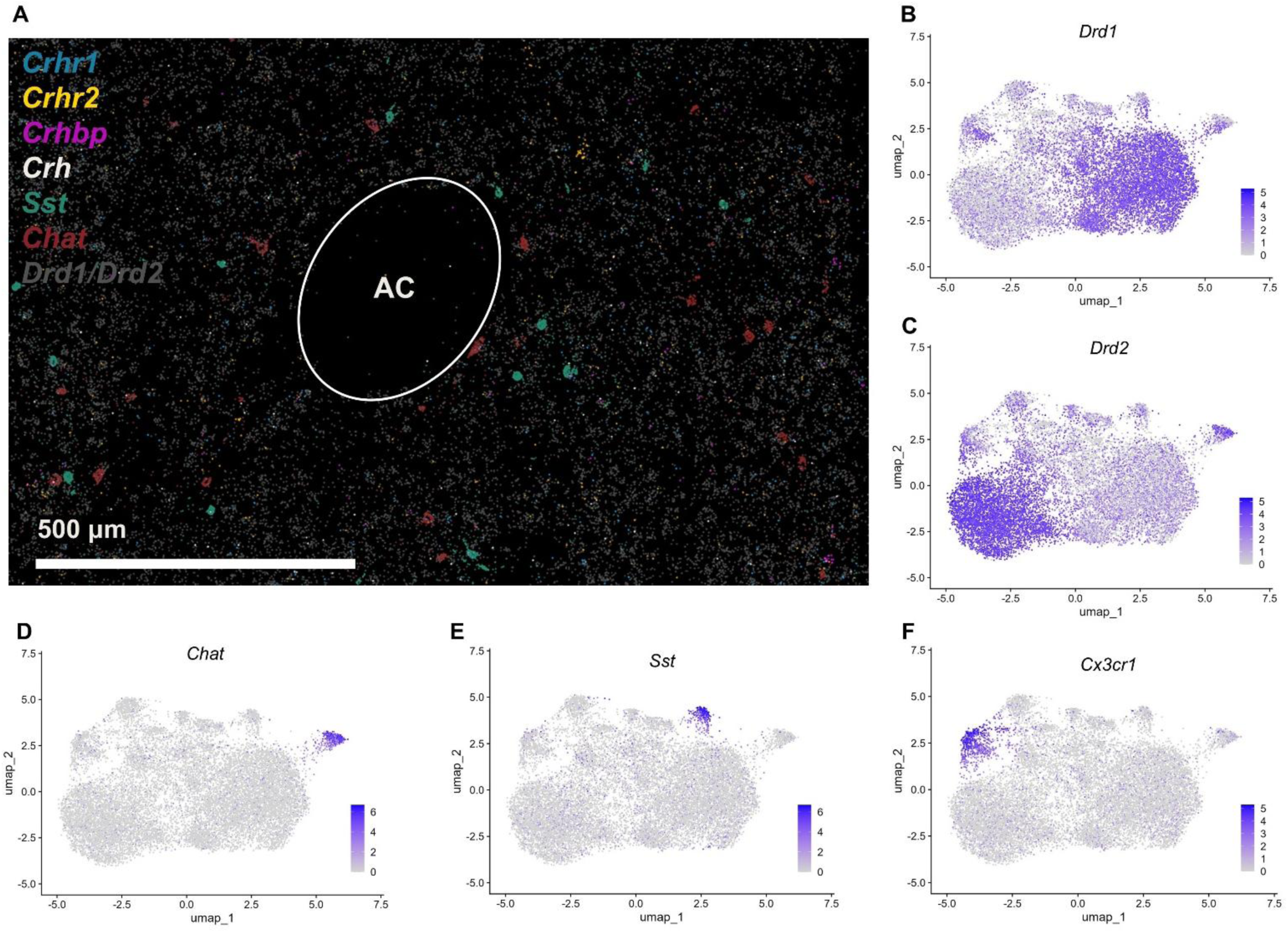
Spatial transcriptomic profiling and UMAP clustering of NAc cell types. **A**, Spatial distribution of genes of interest in a coronal section of the mouse nucleus accumbens. **B** - **F**, UMAP plots of cell type clusters by gene expression.

## Materials and Methods

### Experimental animals

All mice were housed in temperature-controlled and humidity-controlled facilities, with a 12-h light-to-dark cycle and free access to water and standard chow. The ambient temperature was kept at 23–26 °C and humidity was 50–60%. Adult mice aged at least 12 weeks were used for data collection. Males and females were used to assay sex differences in baseline behaviors following early life stress. Only males were used in behavioral assays that manipulated neuropeptide activity and function. The mice used in this study were B6(Cg)-Crh^tm1(cre)Zjh^/J (CRH-ires-CRE) (Jackson Labs, cat. no. 012704), wild-type C57BL/6J (Jackson Labs, cat. no. 000664) and B6.Cg-Gt(ROSA)26Sor^tm14(CAG-tdTomato)Hze^/J mice (Jackson Labs, cat. no. 007914). All experimental procedures were approved by the University of California-Irvine Institutional Animal Care and Use Committee (AUP 24-104 and 21-128) and were in accordance with the guidelines from the National Institutes of Health.

### Limited bedding and nesting (LBN) paradigm

Early-life stress (ELA) was induced on postnatal days (PN) 2 to 9 using our laboratory’s standard protocol (*51, 52*). Conventional mouse cages (Techniplast, cat. no. 1284) were fitted with an aluminum mesh (cat # 4700313244, McNichols, US) platform sitting ∼2.5 cm above the cage floor. Bedding was reduced to cover the cage floor sparsely, and one-half of a single nestlet (5 x 2.5 cm) was provided for nesting material. Control (CTL) dams and pups resided in standard bedding and nesting cages. To control for experimenter handling during the LBN setup, CTL dams and pups were also moved to new cages on PN2. Then, CTL and ELA cages were left undisturbed from PN2 - 9, in a quiet room. On P10, both CTL and ELA groups were transferred to fresh, routine cages.

### Stereotactic viral injection and fiber and/or guide cannula implantation

Mice were anaesthetized with isoflurane (5%). Each mouse was then placed in a robotic stereotaxic apparatus (Stoelting Neurostar StereoDrive, cat. no. 017.005) on a heating pad at 37°C and maintained under isoflurane anaesthesia (1-1.5%) for the duration of the surgery. After shaving the hair and disinfecting the scalp with iodine solution, an incision was made to expose the skull surface. A dental drill (Ideal Micro Drill, Harvard Apparatus, US) was used to perform the craniectomy. Viral infusions were administered using a pulled glass-pipette (Sutter Instruments, cat. no. P97) driven by a Picospritzer III (Parker, US) under nitrogen pressure (35 psi). The total infusion volume was 200 nl for BLA injection and 200 nl for NAc injection. After viral infusion, the pipette was maintained at the injection site for at least 5 min to ensure proper viral distribution and minimize backflow along the injection tract. The viruses were injected at the following coordinates: NAc (anteroposterior (AP), +1.2 mm; mediolateral (ML), 0.5 mm; dorsoventral (DV), 4.4 mm); BLA (anteroposterior (AP), -1.5 mm; mediolateral (ML), 3 mm; dorsoventral (DV), 4.5 mm). For the fiber photometry studies, AAV-hSyn-CRF3.0 or AAV-hSyn-UCN1.0 were injected unilaterally into the left NAc. For the chemogenetic studies, AAV-DIO-hM3D(Gq)-mCherry was injected bilaterally into the BLA. Optical fibers (200 um, 0.5 NA: RWD Life Science, cat. no. R-FOC-BL200C-50NA) were fixed directly above the injection site in the NAc (anteroposterior (AP), +1.2 mm; mediolateral (ML), 0.5 mm; dorsoventral (DV), 4.3 mm). For pharmacological studies, bilateral guide cannulae were fixed 0.5 mm above the target side in the NAc (anteroposterior (AP), +1.2 mm; mediolateral (ML), ±0.75 mm; dorsoventral (DV), 3.8 mm) or unilaterally 0.5 mm above the lateral ventricle (anteroposterior (AP), -0.5 mm; mediolateral (ML), 1 mm; dorsoventral (DV), 1.5 mm). All animals received perioperative pain relief (Buprenorphine 0.1 mg/kg) and recovered from anesthesia on a heating pad before being returned to their home cages for at least 3–4 weeks for viral expression and recovery from surgery before behavioral testing.

### Intra-accumbal or intraventricular infusion

For chemogenetic excitation of CRH+ BLA-NAc inputs, bilateral intra-accumbal infusions of 0.2 µl 1 mM CNO (CNO dihydrochloride, cat. no. HB6149, HelloBio, UK) were delivered via a 30-gauge blunt needle placed 1 mm below the depth of the guide cannula at a constant flow rate of 0.05 µl min^−1^ for four minutes. For CRHR1 antagonism studies in the NAc: 0.2 µl 1 mM α-helicalCRF_9-41_ (MedChemExpress, cat. no. HY-P1294) or 0.2 µl 1 mM NBI 30775 was infused bilaterally into the NAc via a 30-gauge blunt needle placed 1 mm below the depth of the guide cannula at a constant flow rate of 0.05 µl min^−1^ for four minutes. For CRHR1 antagonism studies in the LV: 1 µl 1 mM α-helicalCRF_9-41_ was infused unilaterally into the LV via a 30-gauge blunt needle placed 1 mm below the depth of the guide cannula at a constant flow rate of 0.25 µl min^−1^ for four minutes. For CRH-BP antagonism studies in the NAc: 0.2 µl 1 mM CRF_6-33_ was infused bilaterally into the NAc via a 30-gauge blunt needle placed 1 mm below the depth of the guide cannula at a constant flow rate of 0.05 µl min^−1^ for four minutes. For CRH-BP antagonism studies in the LV: 1 µl 1 mM CRF_6-33_ (MedChemExpress, cat. no. HY-P1297) was delivered via a 30-gauge blunt needle placed 1 mm below the depth of the guide cannula at a constant flow rate of 0.25 µl min^−1^ for four minutes. For CRHR2 antagonism studies in the NAc: 0.2 µl 1 mM antisauvagine30 (MedChemExpress, cat. no. HY-P1107) was infused bilaterally into the NAc via a 30-gauge blunt needle placed 1 mm below the depth of the guide cannula at a constant flow rate of 0.05 µl min^−1^ for four minutes. For CRHR2 stimulation studies in the NAc: 0.2 µl 1 mM Urocortin 2 (MedChemExpress, cat. no. HY-P2847) was infused bilaterally into the NAc via a 30-gauge blunt needle placed 1 mm below the depth of the guide cannula at a constant flow rate of 0.05 µl min^−1^ for four minutes. Following all infusions, the injector remained in the guide cannula for a further four minutes to minimize backflow along the infusion tract. After the conclusion of the infusion, each guide cannula was capped to maintain patency and limit infection.

### Fiber photometry

The fluorescence signal was recorded using a multi-channel fiber photometry device (RWD Life Science Inc., Model: Tricolor Multichannel, cat. no. R821). Fiber photometry data were acquired with 410 and 470 nm µLEDs (200 μm inner core diameter, 0.5 fiber NA (RWD Life Science Inc.), ∼40 µW power from the tip of the connector, Integrating Sphere Photodiode Power Sensor, Thorlabs, cat. no. S140C) at 60 fps (14.7 ms exposure, gain 1.0). Using adjustable time windows (baselines of 20 s or 60 s) in the RWD analysis software, ΔF/F and Z-score data were computed after smoothing (W = 15), baseline correction (Exponential fit, β = 8), and motion correction with the isosbestic signal.

#### Reward cue

Mice were habituated to chocolate cereal (Cocoa Pebbles, Post) for three consecutive days prior to fluorescence recordings. On the test day, the mice were connected to the fiber photometry system and allowed to freely move in the homecage for a minimum of 10 minutes. Following this period, the baseline fluorescence was determined as the mean fitted 470 nm signal within a 20 s time window before the onset of the reward cue, when ∼1 g of chocolate cereal was placed in their home cage. ΔF/F and Z-score data were collected for the following 40 s.

#### Immobilization stress

Mice were naïve to this test. On the test day, the mice were connected to the fiber photometry system and allowed to freely move in the homecage for a minimum of 10 minutes. Following this period, the baseline fluorescence was determined as the mean fitted 470 nm signal within a 20 s time window before the onset of the restraint. ΔF/F and Z-score data were collected for the following 60 s. In additional experiments, the CRHR1 antagonist α-helicalCRF_9-41_ (described above) was infused into the lateral ventricle at least 5 mins prior to the start of restraint stress.

##### CRH-BP (CRF_6-33_)

Mice were naïve to this antagonist. On the test day, the mice were connected to the fiber photometry system and allowed to freely move in the homecage for a minimum of 10 minutes. The baseline fluorescence was determined as the mean fitted 470 nm signal within a 1 min time window before the onset of drug infusion. CRF_6-33_ was infused as described above. ΔF/F and Z-score data were collected from the onset of infusion for 10 min.

### Behavioural assays

All training and testing took place in conventional mouse cages (Techniplast, cat. no. 1284 - home cage), apart from the open field and light/dark transition assays. All testing was carried out during the animal’s active period under low light (< 15 lux). Animal movement in the open field and light/dark transition assays were detected using an automated 3-point software (Ethovision XT15, Noldus, US).

After the completion of behavioral assays, animals were perfused with 4% paraformaldehyde (PFA). The target sites were then examined to verify viral expression and confirm the positions of the optical fibers and/or cannula. Animals that did not exhibit detectable viral expression or had misplaced optical fibers and/or cannula in the target sites were excluded from further analysis.

### Feeding Experimental Device (FED3)

#### Habituation

All mice were habituated to a FED3 device (open ephys, US, cat. no. FED3.1, SKU: OEPS-7510) for 24 hours prior to testing to decrease novelty stress in the homecage environment and increase the probability of pellet collection. During this time, in free-feeding mode, a 20 mg sucrose pellet (5TUT, TestDiet, US, cat. no. 1811149) was automatically dispensed into the feeding well. When the pellet was removed, the time at which it was removed was logged onto an internal microSD card and a new pellet was dispensed. A device was given to each mouse within the first hour of the onset of the dark phase.

#### Testing – Free Feed (Low motivational state)

On the test day, each mouse was given access to the FED3 device in free-feeding mode for 2.5 hours. First, with sucrose pellets for 30 minutes prior to any drug manipulation. After this initial 30-minute test day habituation period, the FED3 device was removed from the homecage, and the mouse was infused with drug for a period of no longer than ten minutes. After the completion of the infusion, the FED3 device was replaced back with the mouse in free-feeding mode and remained undisturbed for 2 hours. The device logged number of pellets collected for the two-hour period following drug infusion.

#### Testing – Progressive Ratio (High motivational state)

On the test day in their homecage, each mouse was given access to the FED3 device coded to a progressive ratio schedule for 12 hours (Richardson and Roberts, 1996). During testing, the nose pokes required to receive a pellet increased exponentially, using Euler’s e, following:

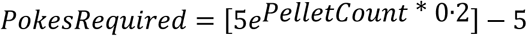

Reward motivation was reported by the number of pokes in the active port, and number of sucrose pellets obtained.

### Open Field

Mice were placed facing the corner of a low lit (< 15 lux measured on arena floor) open field arena (L43 × W43 × H31 cm, Med Associates, Inc, cat. no. ENV-515S-A) and given free access to explore for 10 minutes. The walls inside the arena were white, and the floor was covered in conventional mouse bedding to maintain familiarity with their home cage environment. In analysis, the arena was subdivided into sixteen equal zones (4 inner and 12 outer zones). Center time was measured by collapsing the four inner zones together.

### Light/dark transition

Using a Light/Dark box (Stoelting, US, cat. no. 63101), mice were placed in the brightly lit chamber (> 500 lux) and allowed to freely explore the bright and dark (0 lux) chambers for 10 minutes. Activity (time in bright zone, time in dark zone, latency to zones, and transitions) were tracked using an infrared camera paired with Ethovision XT15 (Noldus, US).

### Immobilization stress

Mice are separated visually and audibly from the cage mates to prevent stress responses in non-recorded mice. Following baseline fiber photometry recordings of at least five minutes, the mouse is placed on a conventional cage grate. Holding the base of the tail, the mouse is held using the nape of the neck. To further immobilize the mouse, it is lifted from the grate, and extended, holding the base of the tail and the nape to ensure no movement. The mouse is held in this position for a total of one minute before being released into their recording cage.

### Immunohistochemistry (IHC)

Mice were transcardially perfused with 4% paraformaldehyde (PFA). Brain tissues were post-fixed in 4% PFA for 4-6 hours at 4°C, and then cryoprotected in a gradient of sucrose solutions (15% and 25%). Coronal sections (20 µm) were prepared and subjected to IHC as described previously (e.g., Chen et al., 2015 Endocrinology; Birnie et al., 2023 NC). Briefly, free-floating sections were treated with 0.3% H_2_O_2_ in 0.01M PBS containing 0.3% Triton X-100 (PBS-T, pH 7.4) for 30 min, and then blocked with 5% normal goat serum (NGS) for 60 min to minimize non-specific binding. After rinsing with PBS-T, sections were incubated for 2 days at 4°C with rabbit anti-RFP antibody (1:2,000, Rockland, cat # 600-401-379). Antibody binding was visualized using Alexa Fluor 488- or 568-conjugated anti-rabbit secondary antibodies (1:400; Invitrogen). Finally, sections were mounted on gelatin-coated slides, air dried, and coverslipped with Slide Mount (Bioenno, Cat#032019 or cat# 032347).

For dual labeling, brain sections were first processed for CRFR1 IHC using a tyramide signal amplification (TSA) approach (Chen et al., 2015). After washes with PBS-T, sections were treated with 0.3% H_2_O_2_ for 30 min, followed by 5% normal donkey serum for 60 min. Sections were rinsed with PBS-T (3 × 5 min) and incubated with goat anti-CRFR1 antibody (1:2,000, aa107-117, N-terminal, Everest Biotech, EB08035, Lot P1 E090910) for 3 days at 4°C. Following PBS-T washes, sections were incubated with HRP-conjugated anti-goat IgG (1:1,000; PerkinElmer) for 1.5 h. Cyanine 3-conjugated tyramide (1:150 in 1× amplification buffer; Akoya) was applied in the dark for 5 min on ice. Next, sections were washed in PBS-T and incubated for 2 days at 4°C with either mouse anti-parvalbumin (1:10,000; Chemicon, MAB354) or mouse anti-ChAT (1:1,000, MA1-25629, clone 7E3-1B8, Affinity BioReagents). Antibody binding was visualized using anti-mouse Alexa Fluor 488 (1:400, Invitrogen, A21206). Finally, sections were mounted and coverslipped using Bioenno Slide Mount (Cat#032019 or cat# 032347).

#### Image acquisition

Confocal images were collected using an LSM-510 confocal microscope (Zeiss) with an apochromatic 10X, 20X, or 63X objective. Virtual z sections of 1 µm were taken at 0.2- to 0.5-µm intervals. Image frame was digitized at 12-bit using a 1024 X 1024 pixel frame size. Z-stack reconstructions and linear adjustments to image brightness and contrast were performed using ImageJ (v2, NIH).

### Multiplexed Error-Robust Fluorescence in situ hybridization (MERFISH)

Two CRH-ires-cre adult male mouse brains (1 CTL, 1 ELA) were analyzed in this study. Mice were single-housed and undisturbed in their home cage for at least 1 week prior to sacrifice. Brains were harvested in the early dark phase (zeitgeber time 1330 – 1500), rapidly frozen in dry-ice chilled 2-methylbutane and embedded in optimal cutting temperature (OCT) compound.

#### Tissue processing

Samples were prepared with a MERscope v1 140 gene panel per manufacturer’s fresh-frozen protocol. Briefly, brains were sectioned in a cryostat at -16°C and mounted directly onto a MERscope slide, post-fixed in 4% paraformaldehyde (PFA), and permeabilized in 70% EtOH for > 24 hours. Tissue was photobleached for six hours during permeabilization. Probe hybridization with a custom 140 gene panel was approximately 40 hours in a humidified 37°C incubator. Each section was run a separate large size slide with an independent study not reported here. Slides were run on a MERscope Ultra machine (software version 234b.241217.1593), housed in the UCI Genomics Research and Technology Hub.

#### Data processing and analysis

Cells were segmented with the MERscope default machine-learning-based tool Cellpose algorithm based on PolyT staining. Cell segmentation quality was visually assessed for alignment to transcript delineated cells. Cells within defined bilateral NAc were virtually dissected from two sections (one per mouse) using Vizgen Vizualizer software (version 2.1.0). NAc cells were exported to a comma-separated value (.csv) file expression matrix for import and analysis in R (version 4.5.3) and Seurat (version 5.4.0). Cells were then filtered to include those with less than 7% MERscope blanks, more than 10 distinct RNA features, and more than forty total RNA counts (*n* = 21,375 cells; *n* = 15,607 post filtered).

Further processing followed standard Seurat workflow including scaling and log-normalization. Datasets from each mouse were integrated using Harmony (version 1.2.4) and then clustered and visualized with uniform manifold approximation and projection (UMAP) and dot plots.

### Micropunch, RNA extraction and qPCR

#### Micropunch

Frozen whole brains were placed in the cryostat (CM3050S, Leica Biosystems, Germany) to increase in temperature to section accurately. Object and chamber temperature were both set at -10°C for sections 200µm thick to prevent tissue cracking. The nucleus accumbens was extracted using the Palkovits Punch technique ^69^. Briefly, NAc coordinates were identified at bregma 1.94mm. Serial sections (5 x 200 µm thick) were taken and ceased at bregma 0.74mm. Bilateral 1mm punches (Uni-Core, Whatman Harris, US) were taken from each of the five NAc sections, stored in a 1.5mL tube (Eppendorf, Germany), and kept on dry ice until ready for RNA extraction.

#### qPCR

*P*unches were processed for total RNA isolation using the Direct-zol RNA purification kit (Zymo Research, Irvine). 100 ng of total RNA was reverse-transcribed using the Transcriptor first-strand cDNA synthesis kit (Roche, IN) with oligo d(T) and random hexamer primers. Each cDNA sample was run in triplicates using FastStart Essential Green Master (Roche, IN) on a Roche Lightcycler 96 system. Data were quantified using the 2^-ΔΔCt method with 18s as housekeeping gene. Primer sequences are provided as follows: 18S: FOR GGGAGCCTGAGAAACGGC; REV GGGTCGGGAGTGGGTAATTT. CRHR1: FOR CTACCAGGGCCCCATGATC; REV CGGACAATGTTGAAGAGAAAGATAAA. CRHR2: FOR CAAGCAGAGAAAGTATGACCTGC; REV GGATACTCCGCAGCACTAGG.

### Statistical analysis

All experiments and data analyses were conducted blind to experimental groups. The number of replicates (*n*) is indicated in figures as lines/dots and in figures legends and refers to the total number of experimental subjects independently treated in each subpanel. Data are presented are mean values, and where applicable, accompanied by SEM. Statistical comparisons were performed using Prism 10 software (GraphPad, USA). Unpaired *t*-tests, paired *t*-tests, and two-way ANOVA with repeated measures were used to test for statistical significance when appropriate. Statistical significance was set at α = 0.05. *P* values are provided in all figures and legends. Experiments were replicated in at least two independent batches, and from at least three different litters. No statistical methods were used to predetermine sample sizes, but our sample sizes are based on those reported in our and others’ publications using similar methods. Individual data points are shown where feasible.

## References

1. Kalmakis, K. A. & Chandler, G. E. Health consequences of adverse childhood experiences: A systematic review. J. Am. Assoc. Nurse Pract. 27, 457–465 (2015).

2. Syed, S. A. & Nemeroff, C. B. Early Life Stress, Mood, and Anxiety Disorders. Chronic Stress (Thousand Oaks, Calif.) 1, (2017).

3. Smith, K. E. & Pollak, S. D. Early life stress and development: potential mechanisms for adverse outcomes. J. Neurodev. Disord. 12, 34 (2020).

4. Taylor, S. E. Mechanisms linking early life stress to adult health outcomes. Proc. Natl. Acad. Sci. 107, 8507–8512 (2010).

5. Teicher, M. H., Samson, J. A., Anderson, C. M. & Ohashi, K. The effects of childhood maltreatment on brain structure, function and connectivity. Nat. Rev. Neurosci. 17, 652–666 (2016).

6. Luby, J. L., Baram, T. Z., Rogers, C. E. & Barch, D. M. Neurodevelopmental Optimization after Early-Life Adversity: Cross-Species Studies to Elucidate Sensitive Periods and Brain Mechanisms to Inform Early Intervention. Trends Neurosci. 43, 744–751 (2020).

7. Birnie, M. T. & Baram, T. Z. The Evolving Neurobiology of Early-life Stress. Neuron (2025).

8. Anda, R. F. et al. The enduring effects of abuse and related adverse experiences in childhood. Eur. Arch. Psychiatry Clin. Neurosci. 256, 174–186 (2006).

9. Green, J. G. et al. Childhood Adversities and Adult Psychiatric Disorders in the National Comorbidity Survey Replication I. Arch. Gen. Psychiatry 67, 113 (2010).

10. Tracy, M., Salo, M., Slopen, N., Udo, T. & Appleton, A. A. Trajectories of childhood adversity and the risk of depression in young adulthood: Results from the Avon Longitudinal Study of Parents and Children. Depress. Anxiety 36, 596–606 (2019).

11. Shonkoff, J. P. et al. The Lifelong Effects of Early Childhood Adversity and Toxic Stress. Pediatrics 129, e232–e246 (2012).

12. LeMoult, J. et al. Meta-analysis: Exposure to Early Life Stress and Risk for Depression in Childhood and Adolescence. J. Am. Acad. Child Adolesc. Psychiatry 59, 842–855 (2020).

13. Heshmati, M. & Russo, S. J. Anhedonia and the Brain Reward Circuitry in Depression. Current Behavioral Neuroscience Reports vol. 2 146–153 (2015).

14. Pizzagalli, D. A. Depression, stress, and anhedonia: toward a synthesis and integrated model. Annu. Rev. Clin. Psychol. 10, 393–423 (2014).

15. Pizzagalli, D. A. Toward a Better Understanding of the Mechanisms and Pathophysiology of Anhedonia: Are We Ready for Translation? Am. J. Psychiatry 179, 458–469 (2022).

16. Nestler, E. J. & Carlezon, W. A. The Mesolimbic Dopamine Reward Circuit in Depression. Biological Psychiatry vol. 59 1151–1159 (2006).

17. Berridge, K. C. & Kringelbach, M. L. Pleasure Systems in the Brain. Neuron vol. 86 646–664 (2015).

18. Stuber, G. D. Neurocircuits for motivation. Science. 382, 394–398 (2023).

19. Floresco, S. B. The Nucleus Accumbens: An Interface Between Cognition, Emotion, and Action. Annu. Rev. Psychol. 66, 25–52 (2015).

20. Herman, J. P. et al. Regulation of the Hypothalamic-Pituitary-Adrenocortical Stress Response. in Comprehensive Physiology vol. 6 603–621 (John Wiley & Sons, Inc., 2016).

21. Kim, J. S., Han, S. Y. & Iremonger, K. J. Stress experience and hormone feedback tune distinct components of hypothalamic CRH neuron activity. Nat. Commun. 10, 5696 (2019).

22. Bale, T. L. & Vale, W. W. CRF and CRF Receptors: Role in Stress Responsivity and Other Behaviors. Annu. Rev. Pharmacol. Toxicol. 44, 525–557 (2004).

23. Roy, A. et al. CSF corticotropin-releasing hormone in depressed patients and normal control subjects. Am. J. Psychiatry 144, 641–645 (1987).

24. Keck, M. E. & Holsboer, F. Hyperactivity of CRH neuronal circuits as a target for therapeutic interventions in affective disorders. Peptides 22, 835–844 (2001).

25. Holsboer, F. Corticotropin-releasing hormone modulators and depression. Curr. Opin. Investig. Drugs 4, 46–50 (2003).

26. Lightman, S. L., Birnie, M. T. & Conway-Campbell, B. L. Dynamics of ACTH and Cortisol Secretion and Implications for Disease. Endocr. Rev. 41, 470–490 (2020).

27. Füzesi, T., Daviu, N., Wamsteeker Cusulin, J. I., Bonin, R. P. & Bains, J. S. Hypothalamic CRH neurons orchestrate complex behaviours after stress. Nat. Commun. 7, 11937 (2016).

28. Chang, S. et al. Tripartite extended amygdala–basal ganglia CRH circuit drives locomotor activation and avoidance behavior. Sci. Adv. 8, (2022).

29. Baumgartner, H. M., Granillo, M., Schulkin, J. & Berridge, K. C. Corticotropin releasing factor (CRF) systems: Promoting cocaine pursuit without distress via incentive motivation. PLoS One 17, e0267345 (2022).

30. Bolton, J. L. et al. Anhedonia Following Early-Life Adversity Involves Aberrant Interaction of Reward and Anxiety Circuits and Is Reversed by Partial Silencing of Amygdala Corticotropin-Releasing Hormone Gene. Biol. Psychiatry 83, 137–147 (2018).

31. Birnie, M. T. et al. Stress-induced plasticity of a CRH/GABA projection disrupts reward behaviors in mice. Nat. Commun. 14, 1088 (2023).

32. Taniguchi, L. et al. Sex and Stress Govern the Function and Innervation of a Basolateral Amygdala to Nucleus Accumbens Corticotropin-Releasing Hormone/GABA-Expressing Projection. J. Neurosci. 46, e1239252025 (2026).

33. Deussing, J. M. & Chen, A. The Corticotropin-Releasing Factor Family: Physiology of the Stress Response. Physiol. Rev. 98, 2225–2286 (2018).

34. Lemos, J. C. et al. Severe stress switches CRF action in the nucleus accumbens from appetitive to aversive. Nature 490, 402–6 (2012).

35. Lemos, J. C., Shin, J. H. & Alvarez, V. A. Striatal Cholinergic Interneurons Are a Novel Target of Corticotropin Releasing Factor. J. Neurosci. 39, 5647–5661 (2019).

36. Lemos, J. C. & Alvarez, V. A. The upside of stress: a mechanism for the positive motivational role of corticotropin releasing factor. Neuropsychopharmacology vol. 45 219–220 (2020).

37. Herman, M. A. et al. Enhanced GABAergic transmission in the central nucleus of the amygdala of genetically selected Marchigian Sardinian rats: Alcohol and CRF effects. Neuropharmacology 67, 337–348 (2013).

38. Cruz, B. et al. Chemogenetic inhibition of central amygdala CRF-expressing neurons decreases alcohol intake but not trauma-related behaviors in a rat model of post-traumatic stress and alcohol use disorder. Mol. Psychiatry 29, 2611–2621 (2024).

39. Itoga, C. A. et al. New viral-genetic mapping uncovers an enrichment of corticotropin-releasing hormone-expressing neuronal inputs to the nucleus accumbens from stress-related brain regions. J. Comp. Neurol. 527, 2474–2487 (2019).

40. Zobel, A. W. et al. Effects of the high-affinity corticotropin-releasing hormone receptor 1 antagonist R121919 in major depression: the first 20 patients treated. J. Psychiatr. Res. 34, 171– 181 (2000).

41. Held, K. et al. Treatment with the CRH1-receptor-antagonist R121919 improves sleep-EEG in patients with depression. J. Psychiatr. Res. 38, 129–136 (2004).

42. Chen, C. & Grigoriadis, D. E. NBI 30775 (R121919), an orally active antagonist of the corticotropin-releasing factor (CRF) type-1 receptor for the treatment of anxiety and depression. Drug Dev. Res. 65, 216–226 (2005).

43. Matikainen-Ankney, B. A. et al. An open-source device for measuring food intake and operant behavior in rodent home-cages. Elife 10, (2021).

44. Rivier, J., Rivier, C. & Vale, W. Synthetic Competitive Antagonists of Corticotropin-Releasing Factor: Effect on ACTH Secretion in the Rat. Science. 224, 889–891 (1984).

45. Wang, H. et al. A tool kit of highly selective and sensitive genetically encoded neuropeptide sensors. Science. 382, (2023).

46. Castro, D. C. & Berridge, K. C. Opioid Hedonic Hotspot in Nucleus Accumbens Shell: Mu, Delta, and Kappa Maps for Enhancement of Sweetness ‘Liking’ and ‘Wanting’. J. Neurosci. 34, 4239– 4250 (2014).

47. Montgomery, K. R. et al. Chemogenetic activation of CRF neurons as a model of chronic stress produces sex-specific physiological and behavioral effects. Neuropsychopharmacology 49, 443– 454 (2024).

48. Seasholtz, A., Valverde, R. & Denver, R. Corticotropin-releasing hormone-binding protein: biochemistry and function from fishes to mammals. J. Endocrinol. 175, 89–97 (2002).

49. Kalin, N. H. Corticotropin-Releasing Hormone Binding Protein: Stress, Psychopathology, and Antidepressant Treatment Response. Am. J. Psychiatry 175, 204–206 (2018).

50. Sutton, S. W. et al. Ligand requirements of the human corticotropin-releasing factor-binding protein. Endocrinology 136, 1097–1102 (1995).

51. Rice, C. J., Sandman, C. A., Lenjavi, M. R. & Baram, T. Z. A Novel Mouse Model for Acute and Long-Lasting Consequences of Early Life Stress. Endocrinology 149, 4892–4900 (2008).

52. Molet, J., Maras, P. M., Avishai-Eliner, S. & Baram, T. Z. Naturalistic rodent models of chronic early-life stress. Dev. Psychobiol. 56, 1675–1688 (2014).

53. Walker, C. et al. Chronic early life stress induced by limited bedding and nesting (LBN) material in rodents: critical considerations of methodology, outcomes and translational potential. Int. J. Biol. Stress 421–448 (2017).

54. Davidson, S. M., Rybka, A. E. & Townsend, P. A. The powerful cardioprotective effects of urocortin and the corticotropin releasing hormone (CRH) family. Biochem. Pharmacol. 77, 141– 150 (2009).

55. Kokkotou, E. et al. Corticotropin-Releasing Hormone Receptor 2-Deficient Mice Have Reduced Intestinal Inflammatory Responses. J. Immunol. 177, 3355–3361 (2006).

56. Flaherty, S. E. et al. Chronic UCN2 treatment desensitizes CRHR2 and improves insulin sensitivity. Nat. Commun. 14, 3953 (2023).

57. Bale, T. L. et al. Mice deficient for corticotropin-releasing hormone receptor-2 display anxiety-like behaviour and are hypersensitive to stress. Nat. Genet. 24, 410–414 (2000).

58. Donner, N. C. et al. Serotonergic systems in the balance: CRHR1 and CRHR2 differentially control stress-induced serotonin synthesis. Psychoneuroendocrinology 63, 178–190 (2016).

59. Skelton, K. H., Owens, M. J. & Nemeroff, C. B. The neurobiology of urocortin. Regul. Pept. 93, 85–92 (2000).

60. Eghbal-Ahmadi, M. et al. The developmental profile of the corticotropin releasing factor receptor (CRF2) in rat brain predicts distinct age-specific functions. Brain Res. Dev. Brain Res. 107, 81– 90 (1998).

61. Eghbal-Ahmadi, M., Avishai-Eliner, S., Hatalski, C. G. & Baram, T. Z. Differential regulation of the expression of corticotropin-releasing factor receptor type 2 (CRF2) in hypothalamus and amygdala of the immature rat by sensory input and food intake. J. Neurosci. 19, 3982–3991 (1999).

62. Chen, K. H., Boettiger, A. N., Moffitt, J. R., Wang, S. & Zhuang, X. Spatially resolved, highly multiplexed RNA profiling in single cells. Science. 348, (2015).

63. Ravichandran, P. et al. Spatiomolecular mapping reveals anatomical organization of heterogeneous cell types in the human nucleus accumbens. Neuron (2026) doi:10.1016/j.neuron.2026.06.002.

64. Rivier, J. et al. Potent and Long-Acting Corticotropin Releasing Factor (CRF) Receptor 2 Selective Peptide Competitive Antagonists. J. Med. Chem. 45, 4737–4747 (2002).

65. Sanson, A., Demarchi, L., Rocaboy, E. & Bosch, O. J. Increased CRF-R1 transmission in the nucleus accumbens shell facilitates maternal neglect in lactating rats and mediates anxiety-like behaviour in a sex-specific manner. Neuropharmacology 265, 110256 (2025).

66. Hupalo, S. et al. Corticotropin-Releasing Factor (CRF) circuit modulation of cognition and motivation. Neurosci. Biobehav. Rev. 103, 50–59 (2019).

67. Heymann, G. et al. Synergy of Distinct Dopamine Projection Populations in Behavioral Reinforcement. Neuron 105, 909–920.e5 (2020).

68. Zalachoras, I. et al. Opposite effects of stress on effortful motivation in high and low anxiety are mediated by CRHR1 in the VTA. Sci. Adv. 8, (2022).

69. Palkovits, M. Isolated removal of hypothalamic or other brain nuclei of the rat. Brain Research vol. 59 (1973).

